# An optimized SP3 sample processing workflow for in-depth and reproducible phosphoproteomics

**DOI:** 10.1101/2025.03.13.643088

**Authors:** Leonard A. Daly, Christopher J. Clarke, Sally O. Oswald, Andris Jankevics, Philip J. Brownridge, Richard A. Scheltema, Claire E. Eyers

**Author notes:** **Corresponding Author. Claire E. Eyers** - Centre for Proteome Research, Institute of Systems, Molecular & Integrative Biology, University of Liverpool, Crown Street, Liverpool L69 7ZB, U.K.; Department of Biochemistry, Cell & Systems Biology, Institute of Systems, Molecular & Integrative Biology, University of Liverpool, Crown Street, Liverpool L69 7ZB, U.K.;, **Leonard A. Daly** - Centre for Proteome Research, Institute of Systems, Molecular & Integrative Biology, University of Liverpool, Crown Street, Liverpool L69 7ZB, U.K.; Department of Biochemistry, Cell & Systems Biology, Institute of Systems, Molecular & Integrative Biology, University of Liverpool, Crown Street, Liverpool L69 7ZB, U.K.

## Abstract

Protein phosphorylation is a ubiquitous post-translational modification (PTM) found across the kingdoms of life, and is critical for the regulation of protein function in health & disease. Advances in high-throughput mass spectrometry have transformed our ability to interrogate the phosphoproteome. However, sample preparation methodologies optimized for phosphoproteomics have not kept pace, compromising the ability to fully exploit these technological advances. In this study, we present an optimized phosphoproteomics workflow using carboxylated SP3 magnetic beads which have simplified proteomics sample preparation. By employing a washing step with 8 M urea and omitting the conventional C_18_ SPE clean-up, we demonstrate a significant improvement in phosphopeptide identifications, with application of this refined protocol to HEK-293T cell extracts increasing the number nearly 2-fold compared to standard SP3 techniques (7908 *cf*. 4129). We also observed a substantial improvement in the detection of multiply phosphorylated peptides. Our findings suggest that the complexity of PTM crosstalk using current peptide-based proteomics workflows is currently under-represented and underscores the necessity of methodological innovations to better capture the intricacies of the phosphoproteome landscape.

## Introduction

Protein phosphorylation is one of the most abundant post-translational modifications (PTMs) with estimates suggesting that ∼90% of human proteins can be phosphorylated *in vivo* at some point in their life cycle^1^. The diverse and critical physiological roles regulated by protein phosphorylation in both a protein-dependent and site-dependent manner means that the phosphoproteome (the phosphorylated protein landscape) has been extensively investigated^2-4^. Continuing advancements in mass spectrometry (MS) and associated (phospho)proteomics analysis techniques have been a cornerstone for interrogation of phos-phorylation-driven signaling networks, with workflows now capable of identifying >10,000 sites in a single experiment^5-9^. As newer technologies (instruments and/or software) become available, a wave of studies follow, each claiming to have greater (phospho)peptide identification rates. However, the advancements afforded by new technologies remain fundamentally dependent on the quality of the samples that are analyzed. Over the past decade, the development of magnetic carboxylated sera-mag speedbeads (SP3 beads) has been a game changer for proteomics pipelines^10, 11^. Their application has facilitated rapid preparation of even the most difficult of samples that have been historically problematic to couple with MS-based workflows in the absence of extensive sample clean-up, which can be both time- and sample-consuming^12-16^. Following commercialization of SP3 beads, a multitude of reports claiming to improve proteome coverage by altering all steps in the sample processing workflow have been published^10, 11, 14, 17-22^.

However, conflicting ‘optimal’ conditions have been reported leading to disparity in the field.

As a specialized branch of proteomics, phosphoproteomics typically requires adapted protocols for optimal sample preparation, consequently few phosphoproteome-specific SP3 studies have been reported^8, 9, 20, 23-26^.

Here, we benchmark SP3 bead technology for phosphoproteomics sample preparation against well-established in-solution processing pipelines, methodically optimizing for improved phosphopeptide coverage. Our final workflow negates the need for the inclusion of protease and phosphatase inhibitors, precipitates in 80% ethanol, includes an additional 8 M urea ‘eluting’ step post-digest and removes the generally accepted C_18_ stagetip clean-up step. This optimized phosphoproteomics pipeline improves both the reproducibility and total phosphopeptide numbers, with nearly a 2-fold improvement in phosphopeptides identified in 2 out of 3 replicates. Moreover, we observe a substantial increase in multiply phosphorylated peptides, with ∼6.7- and ∼42.5-fold more doubly and triply phosphopeptides identified. From a biological standpoint, our findings suggest that current understanding of the phosphoproteome is under-represented with regard to multiply phosphorylated peptides and clustering of phosphorylation sites. Implementation of our new workflow could thus have substantial benefit in advancing understanding of phosphorylation-driven signaling networks, phosphorylation-based cross-talk and total phosphorylation site clustering, particularly when combined with automation and new MS instrumentation and workflows with ever increasing sensitivity.

## Experimental section

### Reagents

Powdered chemical reagents were purchased from Merck and high-performance liquid chromatography (HPLC) grade solvents from ThermoFisher Scientific. All Eppendorf tubes used were ultra-high recovery Eppendorf tubes (STARLAB). Sera-Mag Carboxyl SpeedBeads (SP3) were purchased from Cytivia. Rapigest was purchased from WATERS.

#### Cell culture and sample preparation

Adherent HEK-293T cells were seeded at a density of ∼1.75×10^4^ cells/cm^2^ in DMEM media supplemented with 10% (v/v) fetal calf serum, 1% (v/v) non-essential amino acids and 1% (v/v) penicillin/streptomycin and maintained at 37 ^°^C, 5% CO_2_ until ∼80% confluent (∼5×10^6^ cells/10 cm plate). Cells were washed twice in phosphate buffered saline (PBS) before lysis in 500 μL of stated buffer. All in-solution digest lysis buffers (1% (v/v) Rapigest; 8 M urea; 6 M Guanidine hydrochloride (GuHCl)) were constituted in 100 mM ammonium bicarbonate pH 8 (AmBic). Samples for SP3 processing were lysed using a detergent-based lysis buffer (1% Nonidet P-40, 1% sodium deoxycholate, 1% SDS, 500 mM NaCl, 5 mM EDTA, 100 mM Tris pH 8) ± 1x cOmplete protease inhibitor -EDTA complex (Roche) and phosSTOP (Roche) as stated. Protein content was determined by BCA assay and lysates diluted to 1 μg/μL in relevant lysis buffer prior to reduction and alkylation with dithiothreitol and iodoacetamide^5^. For in-solution digests, samples (250 μg) were diluted 10-fold in 100 mM AmBic pH 8, and Trypsin Gold (Promega) added at a 50:1 (w/w) protein:trypsin ratio and incubated overnight (37 ^°^C with 600 rpm shaking). For SP3 bead-based digests, a 1:1 mixture of magnetic hydrophobic and hydrophilic Seramag-speedbeads (Cytivia) were added at a 1.2:1 (w/w) ratio of beads:protein prior to adding organic solvent (ethanol (EtOH) or acetonitrile (ACN), as stated) to a final concentration of 80% (v/v). Samples were then incubated at 25 ^°^C with 1500 rpm shaking for 30 min. Using a MagRack, the supernatant was discarded and beads washed 3x in 200 μl of 100% of relevant organic solvent. Beads were dried by vacuum centrifugation for 10 min and resuspended in 200 μL of 100 mM AmBic by water bath sonication for 2 min. Trypsin Gold (Promega) was added at a 50:1 (w/w) protein:trypsin ratio, samples were topped up to 220 μL with AmBic and incubated overnight at 37 °C with 1500 rpm shaking. The peptide-containing supernatant was transferred into a fresh tube and, where stated, the remaining beads were washed with a 10:1 (w/v) ratio of beads:stated ‘elution’ solution (300 μg:30 μl, 1% Rapigest, 8 M Urea or 6 M GuHCl, in 100 mM AmBic pH 8) for 30 min with 1500 rpm shaking at 25 °C, before the supernatants were pooled with the corresponding digested material. Trifluoroacetic acid (TFA) was added to a final concentration of 0.5% (v/v) and incubated for 30 min with 1500 rpm shaking at 37 °C followed by 30 min on ice prior to centrifugation (13,000 *g*, 10 min, 4 °C). Clarified supernatants were collected into fresh low-bind tubes, with some being subjected C_18_ stage-tip clean-up. All samples were split 1:99 for analysis of all peptides or titanium dioxide (TiO_2_) enriched phosphopeptide, respectively, prior to vacuum centrifugation or lyophilization to completion, as stated.

### C18 stage-tip clean-up

Samples were subjected to in-house C_18_ stage-tip (Empore Supelco 47 mm C_18_ extraction discs) clean-up prior to LC-MS/MS analysis^27^. Briefly, 3 C_18_ discs were used per 200 μL tip and centrifuged (5000 *g*, 5 min). Stage-tips were equilibrated by sequential washing with 200 μL of methanol, elution solution (75% ACN, 0.1% TFA in water) and wash solution (0.1% TFA in water) prior to loading peptide samples, centrifuging at 4000 *g* for 2 min (or until all liquid had passed through the tip) at room temperature. Flow through was re-applied before washing in 200 μL of wash solution and eluting in 200 μL elution solution. Eluted peptides were dried to completion by vacuum centrifugation.

### Titanium dioxide enrichment

TiO_2_ enrichment was performed as previously described^5^. Briefly, dried peptides were resuspended in TiO_2_ loading solution (80% ACN, 5% TFA, 1M glycolic acid) to a concentration of 1 μg/μl by water bath sonication for 10 min. For post-digest elution optimizations, resuspended peptides were incubated on ice for 30 min prior to centrifugation at 13000 *g* for 10 min at 4 °C and cleared supernatant collected. TiO_2_ resin (GL-Sciences) was added at 5:1 (w/w) TiO_2_ resin:peptide and incubated at 25 ^°^C with 1500 rpm shaking for 30 min before centrifugation at 2000 *g* for 1 min (room temperature) and supernatant removed. TiO_2_ resin-peptide complexes were sequentially washed in 200 μl of TiO_2_ loading solution, solution 1 (80% ACN, 1% TFA in water) and solution 2 (20% ACN, 0.1% TFA in water) prior to vacuum centrifugation for 15 min. Phosphopeptides were eluted in 200 μl of 5% (v/v) ammonium hydroxide in water and shaking at 1500 rpm for 10 min. Samples were centrifuged as before, supernatants collected and dried to completion by vacuum centrifugation.

### Liquid chromatography-tandem mass spectrometry (LC-MS/MS) analysis

All dried peptides were resuspended in MS loading solution (3% ACN, 0.1% TFA in water) by water bath sonication for 10 min, centrifuged (13000 *g*, 4 ^°^C, 10 min) and the cleared supernatant collected. Samples were separated by reversed-phase HPLC over 90 min gradients using an Ultimate 3000 nano system (Dionex)^5^. Data acquisition was performed using a Thermo Orbitrap Fusion Tribrid mass spectrometer (Thermo Scientific) using 32% NCE HCD for 2+ to 5+ charge states over a *m/z* range of 400-1500. MS1 spectra were acquired in the Orbitrap [60K resolution at 200 *m/z*], normalized automatic gain control (AGC) = 50%, maximum injection time = 50 ms and an intensity threshold for fragmentation = 2.5e^4^. For total peptide data, MS2 spectra were acquired in the ion trap set to rapid mode, AGC target = standard, maximum injection time = 35 ms. For phosphopeptide data, MS2 spectra were acquired in the Orbitrap [30K resolution at 200 m/z], AGC target = standard, maximum injection time = dynamic. A dynamic exclusion window was applied to both MS2 approaches for 30 s with a 10 ppm window. Equal volume injections were performed for every experiment.

### Mass spectrometry data analysis

Data was processed with Proteome Discoverer 2.4 (Thermo Scientific) using the MASCOT search engine, searching against the UniProt Human Reviewed database [accessed December 2023]. All data was searched using trypsin (K/R, unless followed by P) with 2 missed cleaves permitted with constant modifications = carbamidomethylation (C) and variable modification = oxidation (M). For phosphopeptide data, the variable modification = phosphorylation (STY) was also included. Label-free quantification was performed using the Minora feature detector node, calculating the area under the curve for *m/z* values. All MS1 mass tolerances = 10 ppm, total peptide MS2 mass tolerance = 0.5 Da, phosphopeptide MS2 mass tolerance

= 0.01 Da. All data were filtered to 1% FDR with the percolator node with standard settings applied. Peptides without a label free quantification measurement in at least 1 replicate of a study were removed from calculations. Non-phosphopeptides were removed from the calculations for phosphopeptide data, unless stated otherwise.

## Results

### SP3 bead-based protocols identify more phosphopeptides with greater consistency than traditional in-solution digest protocols

Multiple publications have reported the benefits of SP3 bead technology to enhance proteomics analysis, primarily due to its versatility and adaptability to different sample lysis conditions, ease of use, and compatibility with automated (robotic) sample processing^10, 11, 14, 17-22^. While a number of these studies have been optimized with respect to total peptide identification numbers, these improvements have typically been modest, and few have considered the effect on phosphopeptide recovery. Primarily these focus on protein:bead ratio^18^, lysis buffer and precipitation conditions^20^ or Eppendorf tube pre-labelling^9^. In agreement with previous studies, our comparison of well-established in-solution sample preparation workflows with the original SP3 protocol employing a detergent-based lysis buffer^10^, demonstrated the benefits of detergent lysis and SP3 processing (Figure 1). This benefit was particularly relevant with respect to Rapigest lysis, given our observations of a >40% increase in identified peptides (≥2 replicates). Interestingly, we also observed notable improvement in the numbers and reproducibility of identification of phosphopeptides using this SP3 workflow, with an increase in phosphopeptide identification of ∼3-fold compared with Rapigest lysis, and over 2-fold compared with either urea or GuHCl. Given these obvious benefits, and the fact that this SP3 workflow was not originally designed with phosphoproteomics in mind, we set out to see if we could further improve this SP3 workflow for phosphopeptide recovery.

**Figure 1:**
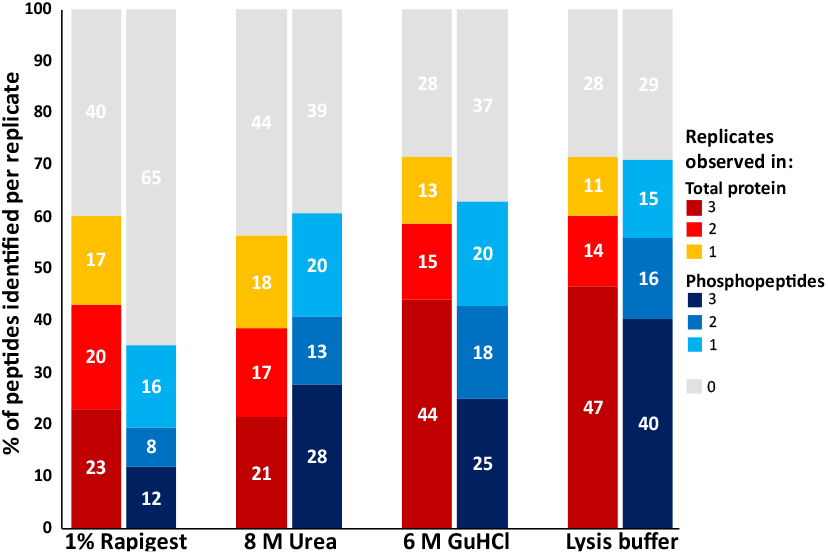
Percentage of total- and phospho-peptides identified using different cell lysis methods. Cells were lysed using the stated condition. Peptides and phospho peptides (after TiO_2_ enrichment) were analyzed by LC-MS/MS. Reported are the numbers of (phospho)peptides identified for each condition (in 1, 2 or 3 replicates (N=3)) as a proportion of the numbers of (phospho)peptides identified across all conditions.

### Ethanol-based precipitation is better for phosphoproteomics

The most controversial step in the SP3 protocol is the precipitation stage: early publications used 50% ACN + 0.1% formic acid^10^, while successive optimizations have suggested that precipitation at neutral pH^14, 17, 20, 22^, ethanol (EtOH)^11, 17, 20, 22^, different concentrations of organic solvent^18, 20, 22^, phosphatase/protease inhibitors^10, 20^, SP3 bead:protein (w/w) ratios^18, 22^ or even bead type^21^ can improve the numbers of peptides identified. However, several groups report conflicting data on their effectiveness.

To simplify workflow optimization, we initially evaluated the ratio of SP3 beads to protein (w/w), and the final concentration of ACN. In agreement with Dagley *et al*^18^, we observed that too high (10:1) a bead:protein (w/w) ratio, or a low final concentration (50%) of ACN decreased the numbers of peptides identified (data not shown). As such, we used a 1.2:1 bead:protein ratio (as previously optimized Dagley *et al*^18^), and 80% ACN as a basis for the next steps. We then evaluated the effect of organic solvent, using EtOH as a replacement for ACN, and a reduction in pH by inclusion of TFA (Figure 2).

**Figure 2:**
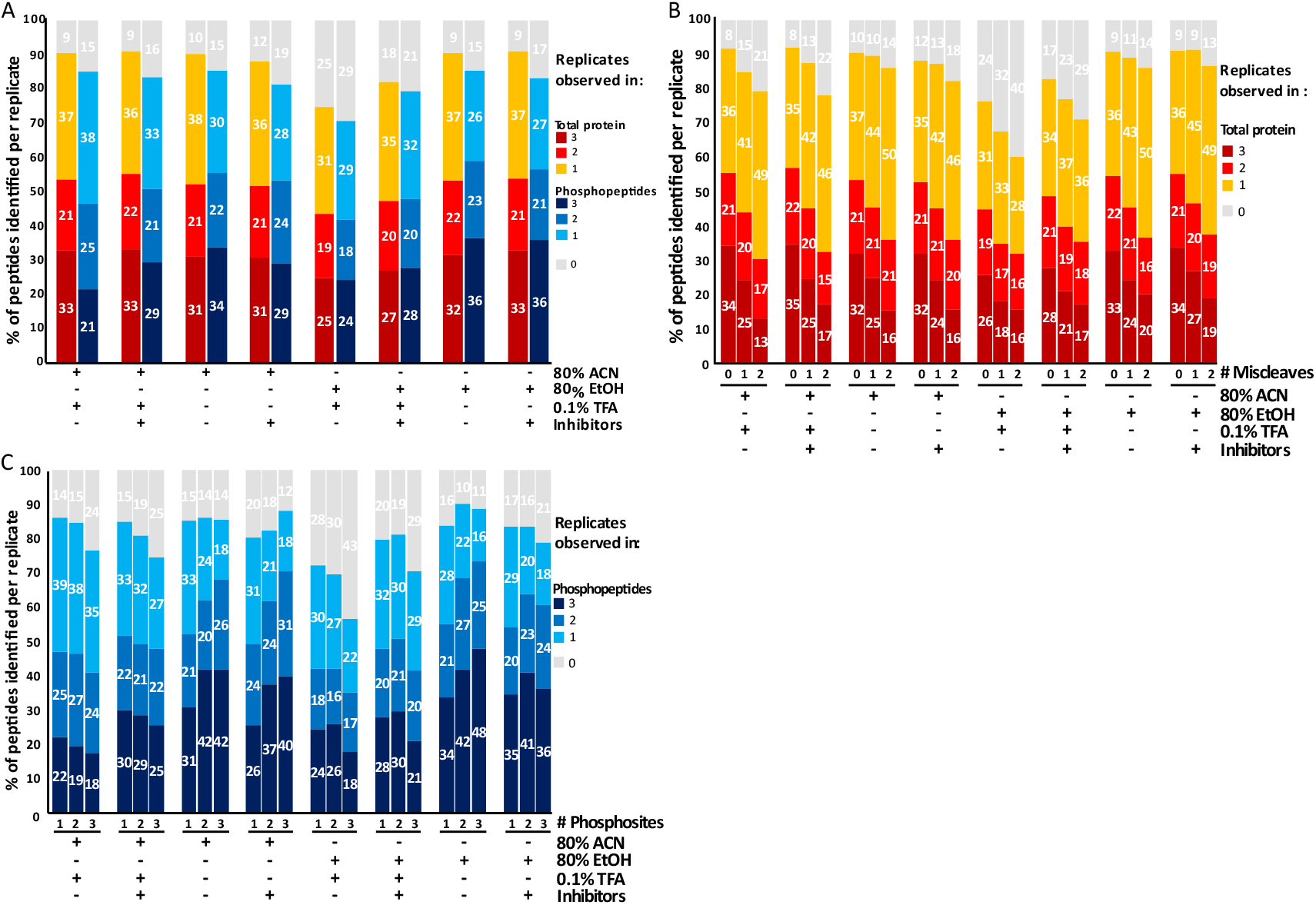
Optimization of SP3 lysis and precipitation conditions. Different precipitation solutions were used as stated for lysates ± complete protease inhibitors and phosSTOP. A total of 17,336 total- and 5313 phospho-peptides were identified and used for normalization respectively. A) Percentage of the total number of total- and phospho-peptides identified across precipitation conditions per replicate. B) Percentage of the total number of peptides identified which contain tryptic miscleaves for total peptide data across precipitation conditions per replicate (0x = 14,668. 1x = 2,398. 2x = 270). C) Percentage of the number of phosphorylation events identified on a peptide across precipitation conditions per replicate (1x = 3,891. 2x = 1,094. 3x = 251).

The number of peptides identified was generally consistent across all conditions, with 52-55% of the 17,336 peptides identified across all conditions being observed in at least 2 replicates of the same condition (Fig 2A). The exception, 80% EtOH + 0.1% TFA, only contained 44% of the peptide complement, supporting observations reported by Moggridge *et al*^*17*^& Leutert *et al*^*20*^. Replacing ACN with EtOH had little discernible effect on the numbers of peptides or proteins identified (Fig 2, Supp. Fig 1). For phosphopeptides, greater variance was observed across the different conditions, with 47-59% (of the 5,313 phosphopeptides identified in total for this experiment, consistent with the expectations of this LC-MS/MS workflow employing a Thermo Fusion Tribrid) being observed (≥2 replicates) dependent on the condition. Notably, a lower pH was generally detrimental, irrespective of whether ACN or EtOH was used (Fig 2A), with exclusion of TFA increasing phosphopeptide numbers by 41%.

Opinions also differ on the benefits of including protease and phosphatase inhibitors during lysis for phosphopeptide workflows. While most pipelines use these as standard^2, 5, 6, 9, 28-33^, Leutert *et al*. reported a notable reduction in the efficiency of digestion, and a >5-fold decrease in peptide identifications in an SP3-enabled pipeline where phosphatase inhibitors (50 mM sodium fluoride, 10 mM sodium pyrophosphate, 50 mM beta-glycerophosphate, 1 mM sodium orthovanadate) and protease inhibitors (protease inhibitor mix, Pierce) were included during lysis^20^. In our hands, inclusion of phosSTOP (Roche) and complete protease inhibitors (Roche) had a limited effect on the number of peptides with a missed cleavage identified, with <5% difference being observed upon inclusion (Fig 2B).

Interestingly, although the presence of inhibitors increased or maintained consistency of identification at the peptide level (irrespective of condition) their inclusion was differentially effective for phosphopeptides, dependent on the pH of the solution (Fig 2A). SP3 sample processing conditions also had a noticeable effect on the number of multiply phosphorylated peptides identified (Fig 2C). For example, there was a ∼20% increase in the number of doubly (2x) and triply (3x) phosphorylated peptides when inhibitors were excluded from the 80% EtOH workflow, with SDS-based lysis conditions that anyway typically render protein kinases inactive^34^. Taken in combination, our data demonstrates that using an SDS-lysis buffer lacking protease/phosphatase inhibitors and precipitating using neutral pH 80% EtOH is optimal for both the number and reproducibility of phosphopeptide identifications, particularly with regard to multiply phosphorylated peptides (Fig 2).

### Removal of SPE clean-up and inclusion of a SP3 ‘elution’ markedly improves phosphopeptide identifications

We have previously observed that standard SP3-based processing of purified α/β-casein tryptic peptides yields fewer phosphopeptides compared to in-solution digests of the same material (data not shown). This supports observations by others that some peptides are not completely eluted from the SP3 bead surface post-digestion in aqueous solutions. Reports suggest that washing in SDS, followed by C_18_ SPE clean-up, can recover an additional 16% median signal intensity^20, 25^. However, the more hydrophilic nature of phosphopeptides^27, 35, 36^ means that they will likely be subject to greater losses during this reversed-phase clean-up step. We therefore wanted to examine the effects (and necessity) of a C_18_ SPE step post SP3 digest, and whether post-digest elution influences the identifiable phosphoproteome. To identify buffers capable of eluting proteins (and presumably peptides) from the surface of SP3 beads, HEK-293T cell lysate precipitated onto SP3 beads was resuspended in a 10:1 (w/v) bead : eluent ratio of either 50 mM AmBic, 1% Rapigest, 8 M urea or 6 M GuHCl, and elutions analyzed by SDS-PAGE alongside an equal protein quantity of non-precipitated lysate (Supp. Fig 2). In-line with Batth *et al*^*25*^, we observed that aqueous solution (50 mM AmBic) was an inefficient protein eluent. However, all other elution conditions performed equally well and were on par with the non-precipitated lysate, confirming efficient elution. Unlike samples in 8 M urea or 6 M GuHCl (which require C_18_ SPE clean-up prior to LC-MS/MS), AmBic and Rapigest samples are LC-MS/MS compatible (post acidification and centrifugal clarification) and can forego this additional sample preparation step. To evaluate the effect and necessity of C_18_ SPE clean-up across elution conditions, we thus analyzed the pooled digest (combined supernatant and elution) by LC-MS/MS with or without C_18_ processing (Fig 3).

**Figure 3:**
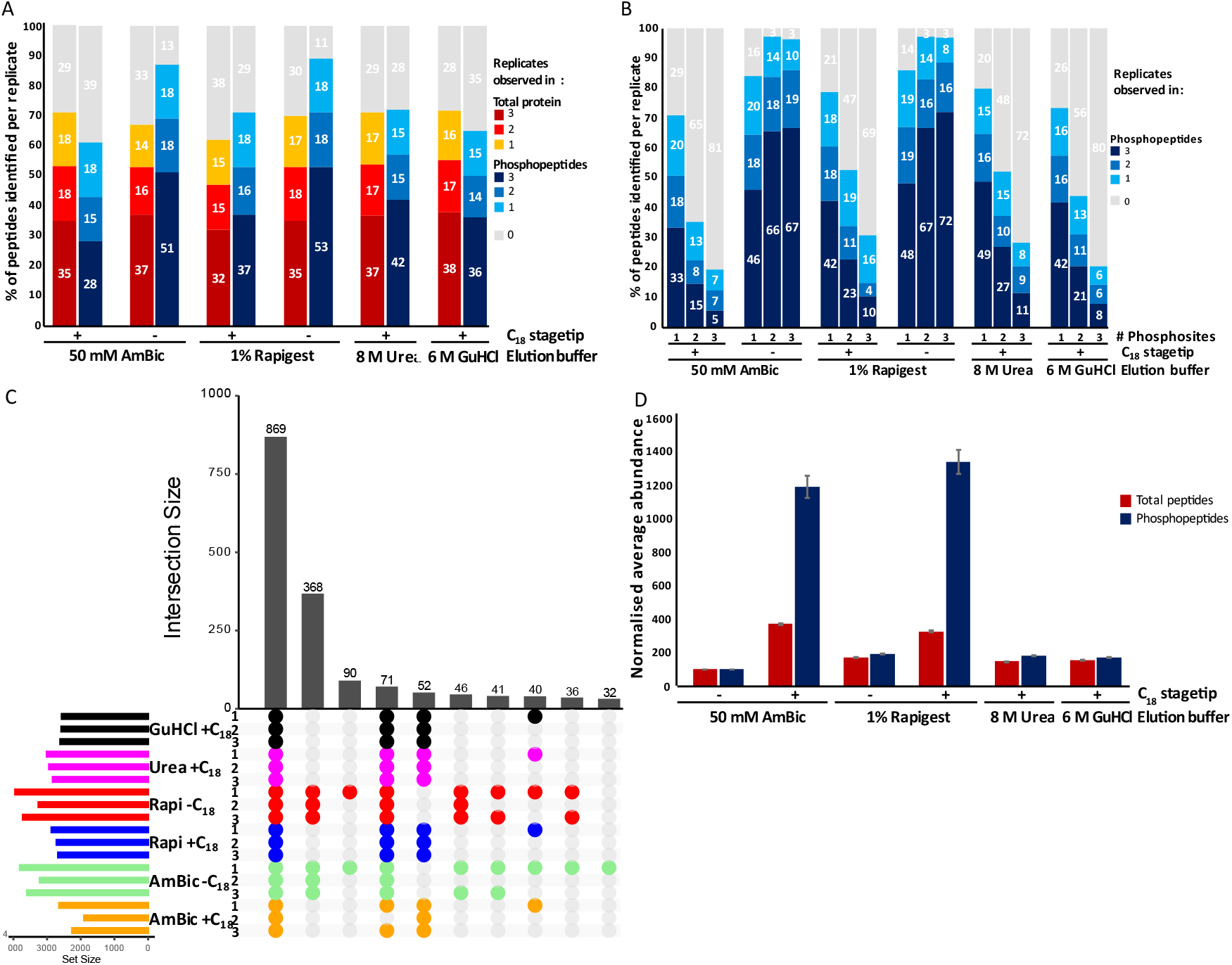
SP3 beads preferentially retain phosphopeptides post digest. Different solutions were used to elute bound peptides from SP3 beads post digest and collection of peptides. A total of 23,817 total- and 5,124 phospho-peptides were identified and used for normalization respectively. A) Percentage of the total number of total- and phospho-peptides identified across different elution solutions per replicate. B) Percentage of the multiply phosphorylated peptides identified across different elution solutions per replicate (1x = 3,844. 2x = 1,051. 3x = 201). C) UpSet plot of phosphopeptides identified across elution solutions and replicates. D) Normalized average abundance of total- and phospho-peptides observed in all replicates of each condition versus 50 mM AmBic +C18 (4,206 and 869 respectively).

At the peptide level, there was little observable difference across any of the elution conditions, with or without C_18_ SPE processing, with ∼50% of all observable peptides being identified in ≥2 replicates (of the ∼23,820 peptides identified across all samples in this experiment, Fig. 3A). Elution conditions and sample clean-up had a much greater effect on phosphopeptide numbers.

Considering C_18_-processed samples in the first instance, 8 M urea elution exhibited the best recovery with ∼31% more phosphopeptides identified (≥2 replicates) versus AmBic processing (Fig 3A). Even greater improvements were observed when C_18_ SPE clean-up was omitted, with >20% more phosphopeptides being identified for AmBic -C_18_ (≥2 replicates) compared to 8 M urea +C_18_ (Fig 3A).

These results were more pronounced for multiply phosphorylated peptides, with ∼1.5, ∼1.8 and ∼2.2-fold increases in singly, doubly and triply phosphorylated peptides, respectively, identified with 8 M urea +C_18_ compared with AmBic +C_18_ (≥2 replicates, Fig 3B). Similarly, omission of C_18_ clean-up from AmBic samples increased observations of singly, doubly and triply phosphorylated peptides by ∼1.3, ∼3.7 and ∼7.3-fold respectively (≥2 replicates). Delving further into the overlap between conditions demonstrates that the biggest cohort of condition-specific phosphopeptide identifications are due to omission of this C_18_ desalting clean-up (Fig 3C), accounting for 368 (∼17%) of observed phosphopeptides across all replicates, a combined improvement of ∼42%.

Finally, label-free quantification (LFQ) confirmed that the increase in phosphopeptide numbers is due to a substantial increase in phosphopeptide recovery (Fig 3D); compared with AmBic +C_18_, there was a ∼2-fold increase in phosphopeptide abundance for all elution +C_18_ SPE treated samples whereas exclusion of the C_18_ SPE step increased the average phosphopeptide abundance by ∼12-fold (equivalent to a 92% loss upon its inclusion, Fig 3D). A similar, but less substantive effect was observed with the non-enriched peptide samples, with omission of the C_18_ clean-up step yielding up to ∼4-fold increase in average abundance of all peptides (Fig 3D).

Given this data, and considering the inexpensive nature of urea, we posited that a phosphoproteomics pipeline that used 8 M urea elution but circumvented C_18_ clean-up could be beneficial, particularly for multiply phosphorylated peptides.

### SP3 digest with urea elution improves identification rates, consistency and enrichment efficiency, particularly for multiply phosphorylated peptides

Although urea-based cell lysis buffers are often used for (phospho)proteomics, they incorporate some form of sample cleanup for salt removal, typically either protein precipitation prior to digestion, or C_18_ SPE (or equivalent) prior to downstream processing or LC-MS analysis^28, 37-39^. Building on our previous findings, we sought to develop a phosphoproteomics workflow that negated the need for such lossy clean-up steps but permitted urea removal prior to phosphopeptide enrichment. Chemically, urea is highly insoluble in ACN^40^, the major constituent of phosphopeptide enrichment buffers (80% ACN in our case), and is volatile. Therefore, we postulated that an evaporative technique (vacuum centrifugation or lyophilization) post digest, followed by ACN-based TiO_2_ phosphopeptide enrichment may overcome any detrimental effects resulting from urea.

We therefore evaluated the utility of either vacuum centrifugation or lyophilization in our SP3 urea elution-based workflow prior to phosphopeptide enrichment. Pooled samples (SP3 digest and urea bead elution) were diluted 4-fold to a final concentration of ∼0.25 M urea, prior to either: i) C_18_ SPE, ii) vacuum centrifugation (as performed for AmBic and Rapigest eluted samples), iii) lyophilization, or iv) 3 rounds of lyophilization (resuspending in 1 ml water between rounds). These four conditions were compared to our previously optimized 50 mM AmBic and 1% Rapigest pipelines (Fig 4).

**Figure 4:**
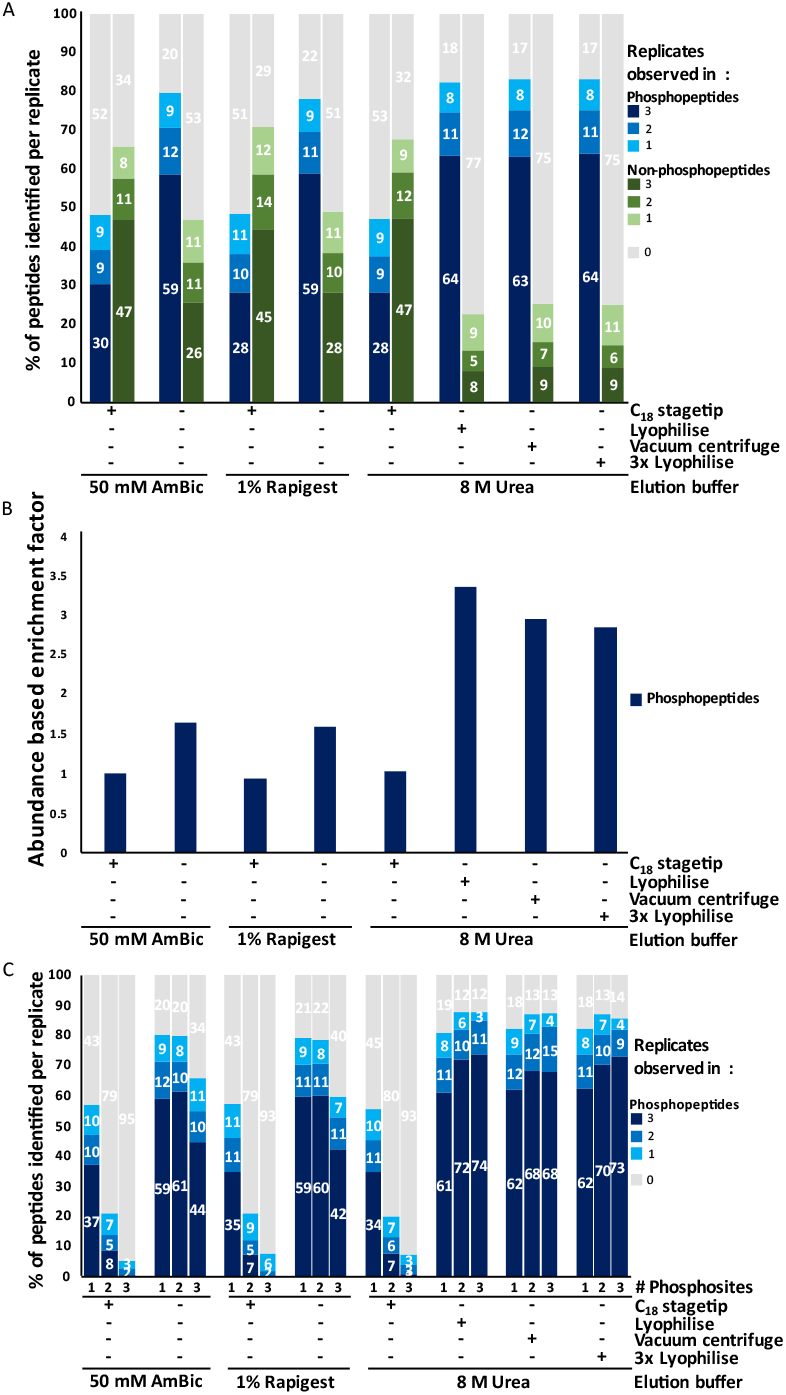
8 M Urea elution post SP3 digest improves phosphopeptide identification rates and TiO_2_ enrichment factors. Different elution solutions and clean-up strategies were used to elute bound peptides from SP3 beads post digest, TiO_2_ data only. A total of 10,508 phospho- and 17,501 non-phospho-peptides were identified and used for normalization respectively. A) Percentage of the total number of phospho- and non-phospho peptides identified post TiO_2_ enrichment across different elution and clean-up strategies per replicate. B) Normalized abundance-based enrichment factor. Average abundance of phosphopeptides observed in all replicates by average abundance of non-phosphopeptides observed in all replicates (1,804 and 682 respectively) normalized against 50 mM AmBic +C_18_. C) Percentage of the multiply phosphorylated peptides identified across different elution solutions per replicate (1x = 8,192. 2x = 1,992. 3x = 299).

Similar to our previous observations, upon removal of the C_18_ SPE clean-up step for the AmBic and Rapigest samples (Fig 3A), all non-SPE-based methods improved phosphopeptide identifications (Fig 4A). Promisingly, all the urea desalting strategies that circumvented C_18_ SPE outperformed our previous ‘best’ methods (AmBic, -C_18_) yielding a 6-8% increase (∼430-490 peptides) in the numbers of phosphopeptides identified (≥2 replicates, Fig 4A), with no substantial difference being observed dependent on the evaporative process used (Supp Fig 3A). While there were slight gains in the numbers of multiply phosphorylated peptides with a single lyophilization step (Fig 4C), it is worth noting that vacuum centrifugation benefits from speed.

An additional benefit of evaporative urea removal was the improved TiO_2_-based phosphopeptide enrichment. Phosphopeptides accounted for only ∼30% of the total peptides identified using the ‘standard’ AmBic +C_18_ TiO_2_ workflow (10,069 non-phosphorylated peptides *cf* 4,129 phosphopeptides, observed in ≥2 replicates), increasing to >50% upon removal of C_18_ cleanup (6,319 non-phosphorylated peptides *cf* 7,418 phosphopeptides, observed in ≥2 replicates). Using urea + lyophilization improved phosphopeptide enrichment over 3-fold compared with the standard workflow (2,345 non-phosphorylated peptides *cf* 7,850 phosphopeptides, observed in ≥2 replicates *i*.*e*. 77% enrichment; Fig 4A/B). UpSet plots and LFQ analysis (Supp. Fig 3) further highlight the efficiency of our urea elution strategy, revealing a bias of C_18_ SPE for preferential selection of non-phosphopeptides (Supp. Fig 3A/B). Indeed, the relative phosphopeptide abundance increases by >18-fold for those workflows that do not include a C_18_ SPE step (equivalent to a ∼94% loss).

Finally, and in-line with our observations with both AmBic and Rapigest in the absence of C_18_ SPE (Fig 3D), we observed substantial improvement in the identification of multiply phosphorylated peptides, in particular for the three urea-based evaporative workflows. All three were similarly beneficial, resulting in ∼1.6-, ∼6- and ∼36-fold increases in singly, doubly and triply phosphorylated peptides respectively (≥2 replicates), compared with the standard AmBic +C_18_ method (Fig 4C). Overall, our updated sample preparation workflow demonstrates substantive benefits for phosphopeptide analysis, improving sensitivity, reproducibility, throughput and sustainability given the faster workflow and reduced consumables requirements.

## Conclusions

In considering the disparate published pipelines for (phospho)proteomics sample processing, we evaluated and optimized in-solution and SP3-based digestion workflows, focusing on the number of (phospho)peptides identified, and the reproducibility of identification.

As previously reported, we observed (marginally) better data firstly when cells were lysed in an SDS-based lysis buffer lacking protease and phosphatase inhibitors, and secondly when ethanol rather than ACN was used for SP3 precipitation. However, unlike previous studies, we did not observe any notable increase in the rate of trypsin missed cleavages when inhibitors were omitted, and we optimally used a higher concentration of ethanol ^11, 17, 20^. We therefore recommend that for these types of cell-based (phospho)proteomics investigations that inhibitors be excluded for the sake of data quality and cost. However, it is worth noting that the levels of cellular proteases are much higher in some other biological/tissue samples (e.g. pancreas, liver) and as such their omission should be re-evaluated in context.

The single biggest improvement that we identified was removal of the routine C_18_ SPE stage-tip sample clean up step ^17, 18, 20, 23-25^, which resulted in a dramatic ∼12-fold increase in phospho-peptide abundance and ∼1.6-fold more phosphopeptides being identified, with relatively small differences in the numbers of non-phosphorylated peptides. We hypothesize that this is due to the increased hydrophilicity of phosphopeptides *cf* non-phosphorylated peptides ^27, 35, 36^, hence loss of these are preferentially exacerbated during this SPE step.

The second biggest improvement was post-digest washing of SP3 beads in 8 M urea. Interestingly, our data shows minimal benefit for total peptide identifications but increased the identified phosphoproteome >1.3-fold, suggesting preferential binding of these typically more hydrophilic and acidic peptides to these carboxylated SP3 beads. The final protocol, which included evaporative removal (lyophilization or vacuum centrifugation) and precipitation of the urea eluent post-digest, prior to TiO_2_ phosphopeptide enrichment increased phosphopeptide identification rates by nearly 2-fold (≥2 replicates) compared with current ‘standard’ SP3 protocols. This substantial improvement in phosphopeptide identifications is likely in part due to the greater phosphopeptide enrichment afforded with this urea-based workflow.

Overall, our data shows that SP3 bead elution with 8 M urea followed by non-C_18_-based salt removal substantially improves phosphopeptide recovery and thus the reproducibility and numbers identified. The amendments described here are readily adaptable to robotics coupling, which is a major advantage of SP3 technology. Biologically, our data undoubtably raises questions regarding our current understanding of proximal phosphorylation crosstalk, given the marked increase in the identification of doubly and triply phosphorylated peptides, equivalent to >6- and >36-fold more versus typical SP3 protocols. Additionally, while not considered here, it is important to state that our optimized pipeline (up to TiO_2_ enrichment) is pH neutral and performed at room temperature, and is therefore compatible with the preparation of samples for the analysis of atypical sites of phosphorylation (His, Lys, Arg, Cys, Asp, Glu^28, 41, 42^) and may be relevant for the analysis of other acidic PTMs including sulfation^27^.

## ASSOCIATED CONTENT

## AUTHOR INFORMATION

### Author Contributions

L.A.D. performed all sample preparation, enrichment and mass spectrometry analysis, contributed to design of experiments and manuscript writing. C.J.C. and S.O.O. contributed to sample preparation. A.J. and R.A.S. supported data analysis pipelines. P.J.B. supported mass spectrometry methods development. C.E.E. contributed to design of experiments, analysis of data, and manuscript writing. The manuscript was written by L.A.D. and C.E.E. All authors have given approval to the final version of the manuscript.

### Notes

The authors declare no competing financial interest.

All MS data has been deposited at the ProteomeXchange Consortium (http://proteomecentral.proteomexchange.org) via the PRIDE partner repository with the data set identifier PXD061013. **Username:**. **Password:** j44E8kW6v9iU.

## ACKNOWLEDGMENTS

This work is supported by funding from the Biotechnology and Biosciences Research Council (BBSRC; BB/S018514/1, BB/M012557/1, BB/S017054/1, BB/R000182/1, BB/X002780/1 and BB/W00349X/1) and the Royal Society Wolfson Foundation (RSWF\R2\232006).

---

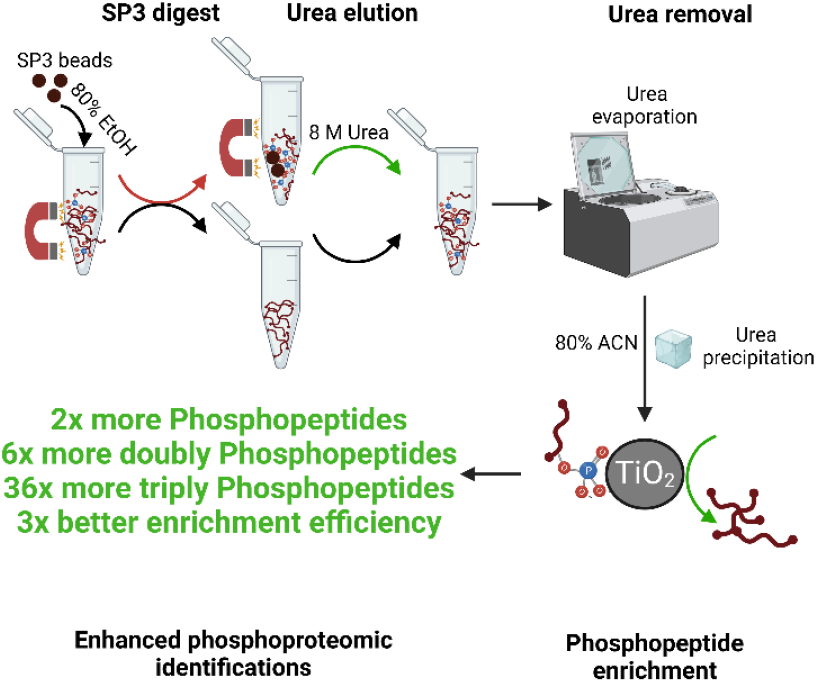
Table of Contents

## Notes

### Competing Interest Statement

The authors have declared no competing interest.

