## Supplementary material for "An optimized SP3 sample processing workflow for in-depth and reproducible phosphoproteomics": Supp. Figure 1

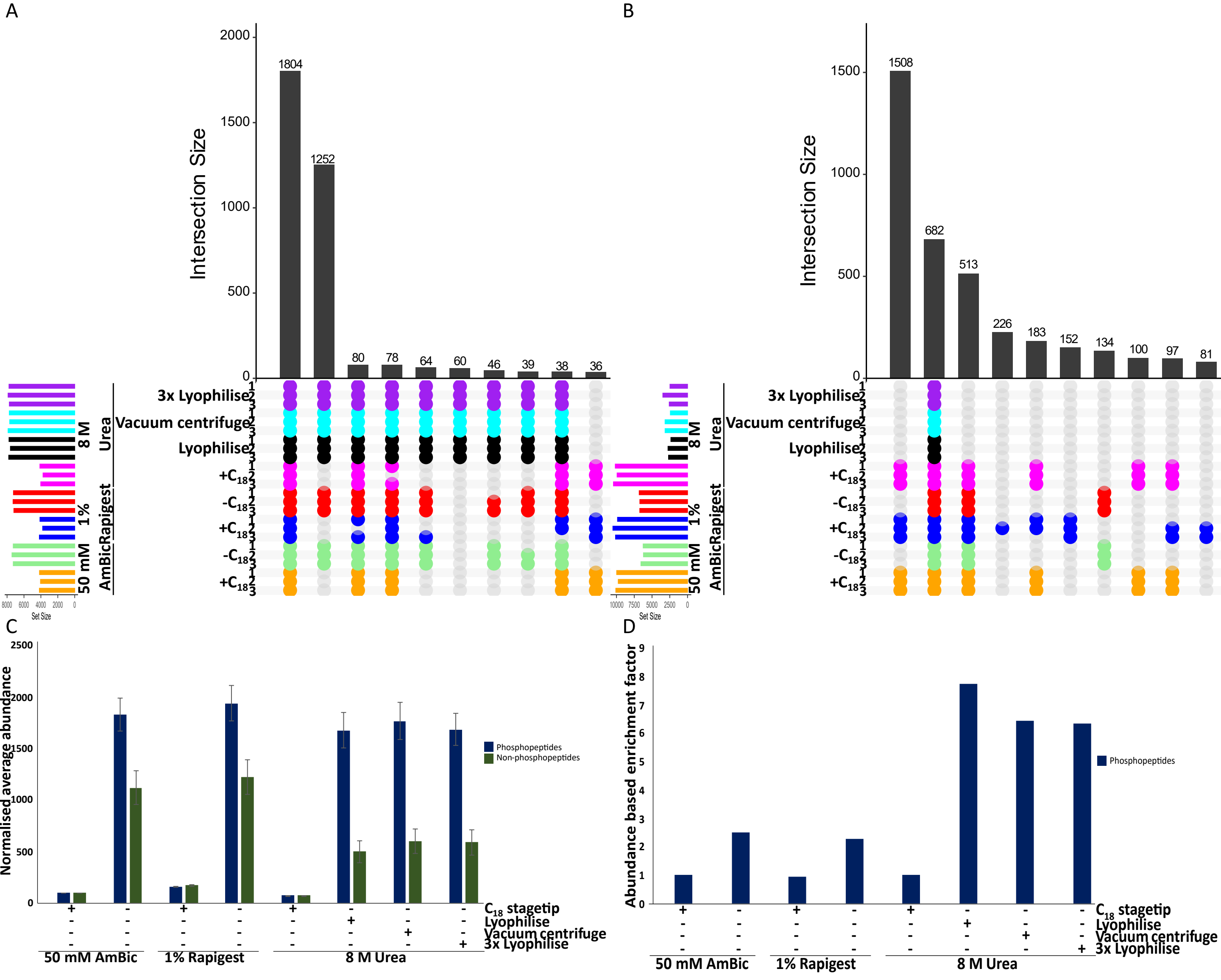


***Supp. Figure 3: Evaporative removal for urea elution strategies increase phosphopeptide identifications and enrichment efficiency.*** *SP3 beads were post digest washed in stated solution, and subject to either ±C18 SPE clean-up or, for urea removal, evaporative strategies prior to $TiO_2$ phosphopeptide enrichment. A total of 10,508 phospho- and 17,501 non-phospho- peptides were identified post enrichment. A) UpSetPlot of phosphopeptides identified across conditions. B) UpSetPlot of non-phosphopeptides identified across conditions. C) Normalized average abundance of phospho- and non-phospho- peptides observed in all replicates of each condition versus 50 mM AmBiC +C18 (1804 and 682 respectively). D) Abundance based enrichment factor. Calculation (Average abundance of phosphopeptides / average abundance of non-phosphopeptides) post removal of the Top 20 differentially abundant non-phosphopeptides.*
