## Supplementary material for "An optimized SP3 sample processing workflow for in-depth and reproducible phosphoproteomics": Supp Figure 2

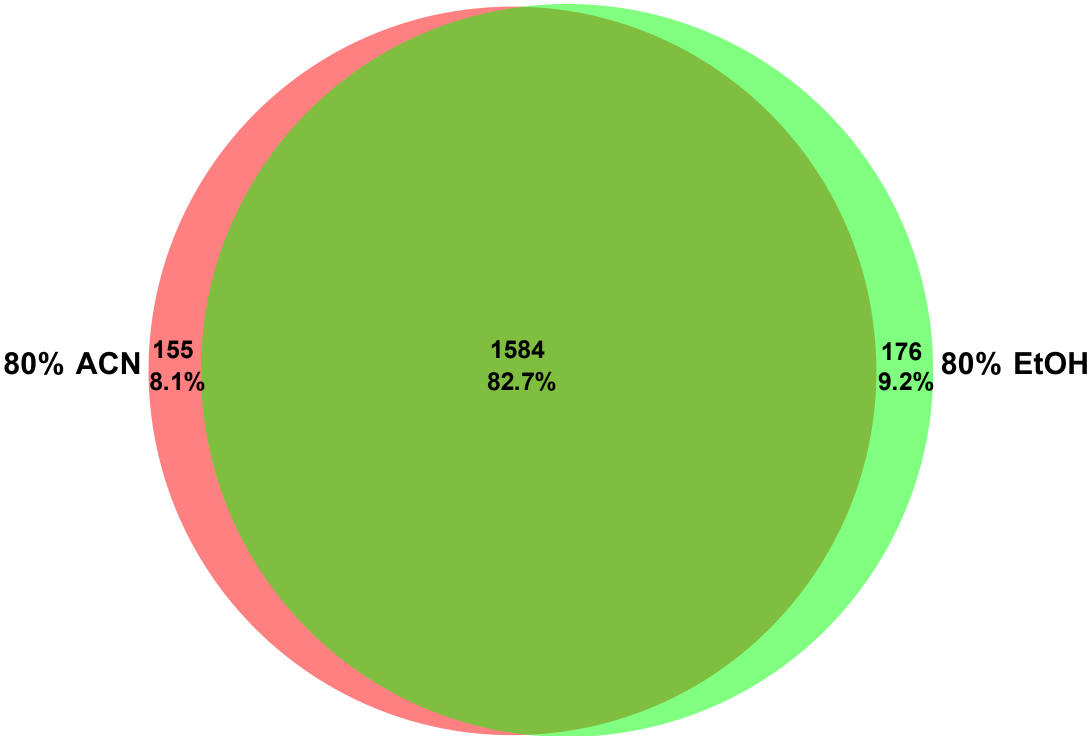


***Supp. Figure 1: Protein level differences between precipitation solvents.*** *Lysate without protease inhibitors and phosSTOP were precipitated with different organic solvents. Venn diagram of protein Identifications observed in ≥2 replicates.*
