## Supplementary material for "An optimized SP3 sample processing workflow for in-depth and reproducible phosphoproteomics": Supp Figure 3

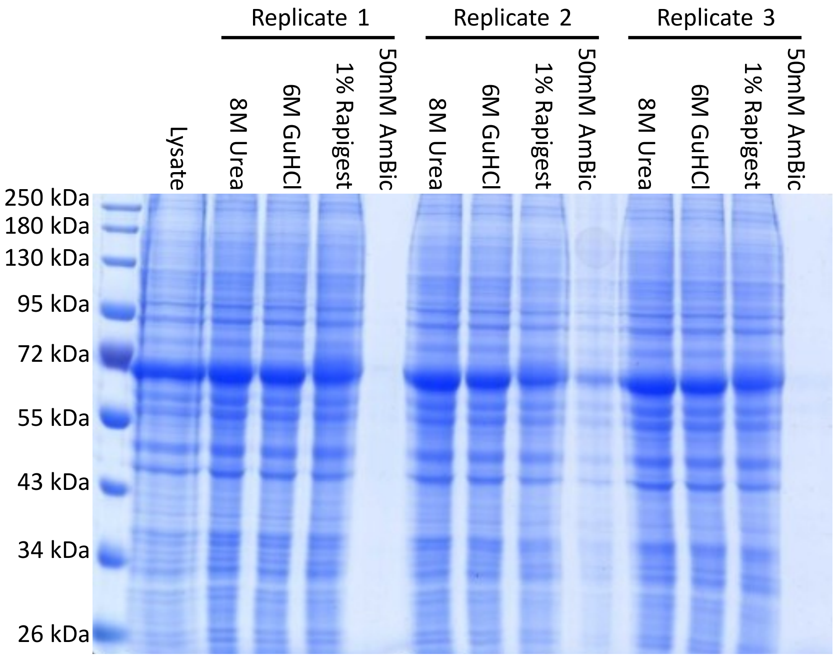


***Supp. Figure 2: Protein elution strategies from SP3 beads****Coomassie gel of lysate (without protease inhibitors and phosSTOP) precipitated with 80% EtOH onto SP3 beads and eluted using stated solution, alongside an equal loading of non-precipitated lysate.*
